# Molecular characterization of the coat protein gene revealed considerable diversity of viral species complex in Garlic (*Allium sativum* L.)

**DOI:** 10.1101/2020.12.03.409680

**Authors:** Abel Debebe Mitiku, Dawit Tesfaye Degefu, Adane Abraham, Desta Mejan, Pauline Asami, Solomon Maina, Timothy Holton

## Abstract

Garlic is one of the most crucial Allium vegetables used as seasoning of foods. It has a lot of benefits from the medicinal and nutritional point of view; however, its production is highly constrained by both biotic and abiotic challenges. Among these, viral infections are the most prevalent factors affecting crop productivity around the globe. This experiment was conducted on eleven selected garlic accessions and three improved varieties collected from different garlic growing agro-climatic regions of Ethiopia. This study aimed to identify and characterize the isolated garlic virus using the coat protein (CP) gene and further determine their phylogenetic relatedness. RNA was extracted from fresh young leaves, thirteen days old seedlings, which showed yellowing, mosaic, and stunting symptoms. Pairwise molecular diversity for CP nucleotide and amino acid sequences were calculated using MEGA5. Maximum Likelihood tree of CP nucleotide sequence data of *Allexivirus* and *Potyvirus* were conducted using PhyML, while a neighbor-joining tree was constructed for the amino acid sequence data using MEGA5. From the result, five garlic viruses were identified viz. *Garlic virus C* (78.6 %), *Garlic virus D* (64.3 %), *Garlic virus X* (78.6 %), *Onion yellow dwarf virus* (OYDV) (100%), and *Leek yellow stripe virus* (LYSV) (78.6 %). The study revealed the presence of complex mixtures of viruses with 42.9 % of the samples had co-infected with a species complex of *Garlic virus C, Garlic virus D, Garlic virus X,* OYDV, and LYSV. Pairwise comparisons of the isolated *Potyviruses* and *Allexiviruses* species revealed high identity with that of the known members of their respected species. As an exception, less within species identity was observed among *Garlic virus C* isolates as compared with that of the known members of the species. Finally, our results highlighted the need for stepping up a working framework to establish virus-free garlic planting material exchange in the country which could result in the reduction of viral gene flow across the country.

**Author Summary:** Garlic viruses are the most devastating disease since garlic is the most vulnerable crop due to their vegetative nature of propagation. Currently, the garlic viruses are the aforementioned production constraint in Ethiopia. However, so far very little is known on the identification, diversity, and dissemination of garlic infecting viruses in the country. Here we explore the prevalence, genetic diversity, and the presence of mixed infection of garlic viruses in Ethiopia using next generation sequencing platform. Analysis of nucleotide and amino acid sequences of coat protein genes from infected samples revealed the association of three species from *Allexivirus* and two species from *Potyvirus* in a complex mixture. Ultimately the article concludes there is high time to set up a working framework to establish garlic free planting material exchange platform which could result in a reduction of viral gene flow across the country.

## Introduction

Garlic (*Allium sativum* L.) belongs to the family *Alliaceae* and genus *Allium*. It is one of the main Allium known and distributed throughout the world, and the second most widely used next to onion [1]. Garlic is the most ancient cultivated herbs grown for culinary, nutritional and medical purposes [2, 3]. Its flavoring nature makes it one of the top priority vegetables used as a spice for foods [4]. Albeit garlic is one of the most crucial vegetable crops, its production, and productivity is highly constrained by biotic and abiotic challenges starting from the seedling stage to harvest.

Viral diseases are the most important biotic constrain that determine the yield and quality of garlic production in the world [5]. Twelve major viruses which are found in three genera (*Potyvirus, Allexivirus*, and *Carlavirus*) are the most common viral species consistently found in garlic infection. They are *Onion yellow dwarf virus* (OYDV), *Leek yellow stripe virus* (LYSV), *Garlic virus A* (Gar-VA), *Garlic virus B* (Gar-VB), *Garlic virus C* (Gar-VC), *Garlic virus D* (Gar-VD), *Garlic virus E* (Gar-VE) and *Garlic virus X* (Gar-VX), *Garlic mite born filamentous viruses* (G-Mb Fv) and *Shallot virus X* (Sh-V X), *Garlic common latent virus* (GCLV) and *Shallot latent viruses* (SLV) [6–9].

Enzyme-linked immunosorbent assays (ELISA) and polymerase chain reactions (PCR) have been playing a key role in easier identification of viruses although they have many drawbacks. For instance, the need for specific assay for each pathogen hosts interaction and limited ability to differentiate Allium viruses because of multiple infections [10–12]. On the contrary, the advent of modern techniques viz. Next-Generation Sequencing revolutionized the detection, discovery, and identification of viruses [13–15].

Research result in many countries showed that viral isolates from the different genera either in single or complex had been detected. A species complex of *Onion yellow dwarf virus* (OYDV) and *Leek yellow stripe virus* (LYSV) were the most crucial viruses in Italy, Greek, and Brazil [9, 16–18]. Whereas *Garlic virus A* (Gar-VA), *Garlic virus B* (Gar-VB), *Garlic virus C* (Gar-VC), *Garlic virus D* (Gar-VD), *Garlic virus E* (Gar-VE) and *Garlic virus X* (Gar-VX) were the most prevalent species from *Allexivirus* [7, 19]. It is not uncommon to see viral species complex from the three genera as reported by different authors [5, 20–23].

In Ethiopia, as in other parts of the world, garlic is grown for different purposes mainly as a spice for seasoning of foods. Like many other countries in the world, garlic infecting viruses are the aforementioned production constraint in the country. In this regard, insect pests play significant impact through vectoring the viruses throughout the garlic production belts of Ethiopia [24, 25]. Furthermore, almost all garlic production systems of Ethiopia used bulbs as a planting material which raises a concern about the presence of considerable diversity of garlic viral species complex. However, so far very little is known on identification, diversity and dissemination of garlic infecting viruses in the country [26–29]. In order to address this problem, a comprehensive study to evaluate the prevalence and characterization of the viruses existing at the moment in the country is inevitable. Hence, this study was designed to identify and characterize the genetic variability and phylogenetic relatedness among the detected virus species using the coat protein gene.

## Materials and Method

### Plant material and filed screening

Garlic cloves of eleven accessions and three improved varieties collected from major garlic growing agro-climatic regions of Ethiopia by Debre-Zeit Agricultural Research Centre were used as experimental material (Table 1)

**Table 1.**
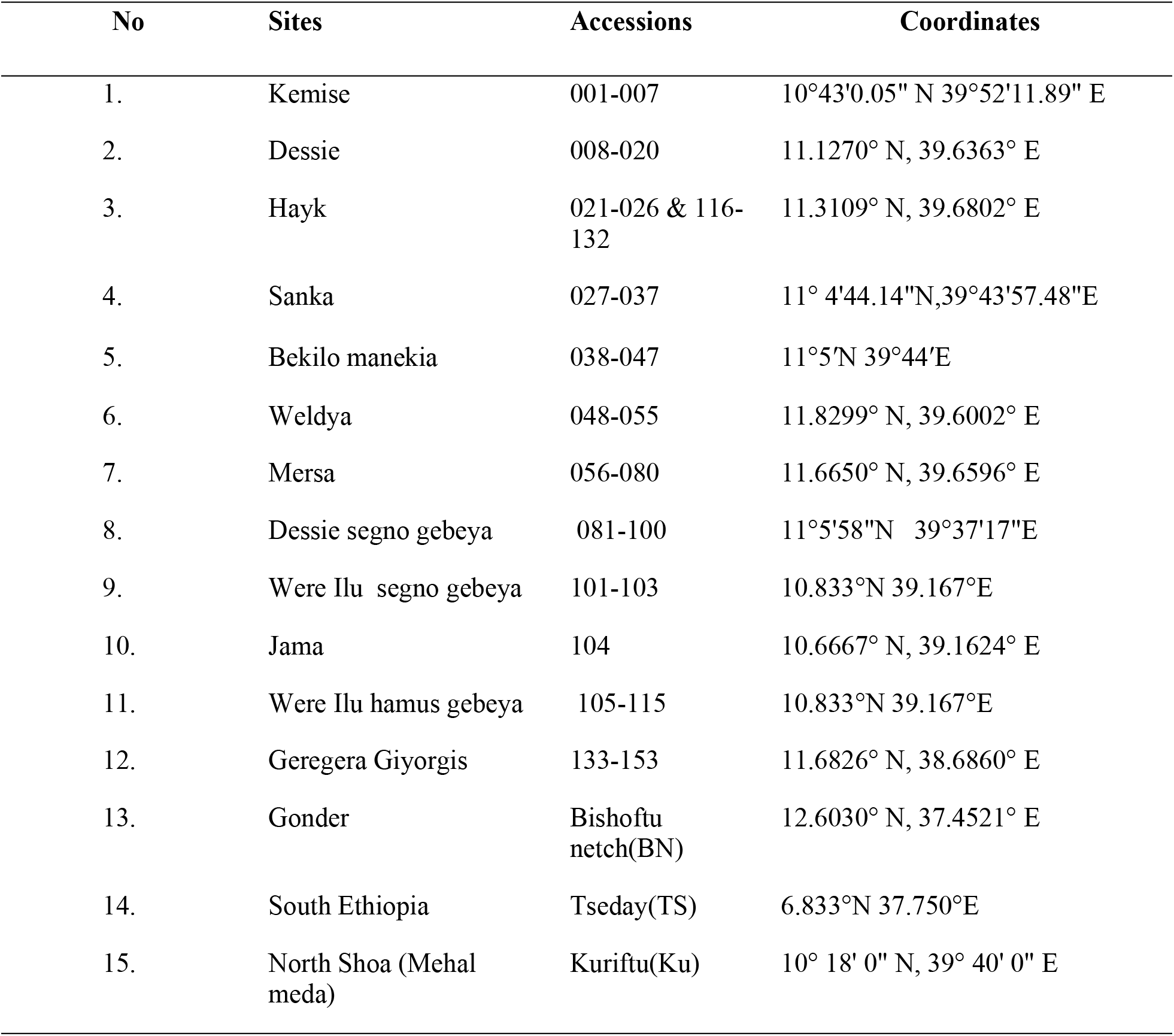
Sample collection sites in Ethiopia

| No | Sites | Accessions | Coordinates |
| --- | --- | --- | --- |
| 1. | Kemise | 001-007 | 10°43'0.05" N 39°52'11.89" E |
| 2. | Dessie | 008-020 | 11.1270° N, 39.6363° E |
| 3. | Hayk | 021-026 & 116-132 | 11.3109° N, 39.6802° E |
| 4. | Sanka | 027-037 | 11° 4'44.14"N,39°43'57.48"E |
| 5. | Bekilo manekia | 038-047 | 11°5'N 39°44'E |
| 6. | Weldya | 048-055 | 11.8299° N, 39.6002° E |
| 7. | Mersa | 056-080 | 11.6650° N, 39.6596° E |
| 8. | Dessie segno gebeya | 081-100 | 11°5'58"N 39°37'17"E |
| 9. | Were Ilu segno gebeya | 101-103 | 10.833°N 39.167°E |
| 10. | Jama | 104 | 10.6667° N, 39.1624° E |
| 11. | Were Ilu hamus gebeya | 105-115 | 10.833°N 39.167°E |
| 12. | Geregera Giyorgis | 133-153 | 11.6826° N, 38.6860° E |
| 13. | Gonder | Bishoftu netch(BN) | 12.6030° N, 37.4521° E |
| 14. | South Ethiopia | Tseday(TS) | 6.833°N 37.750°E |
| 15. | North Shoa (Mehal meda) | Kuriftu(Ku) | 10° 18' 0" N, 39° 40' 0" E |

### RNA extraction

All the garlic samples were planted in a net house at Bioscience Eastern and Central Africa-International Livestock Research Institute (BecA-ILRI) Hub, Nairobi in 2014. Then fresh young leaves which showed yellowing, mosaic and stunting symptoms from thirteen days old potted seedlings were collected to extract RNA using ZR plant RNA Miniprep kits (zymoreserch.com) following the manufacturer’s instruction. RNA quantity and quality were evaluated using NanoDrop Spectrophotometer.

### cDNA library preparation and sequencing

Ribozero RNA-Seq library was prepared following Illumina TruSeq RNA sample preparation Kits (www.Illumina.com/TruSeq). After purifying the mRNA, they were fragmented into small pieces and copied into first-strand cDNA templates using reverse transcriptase random primer followed by synthesis of the second strand using DNA polymerase I. Ends of each fragment were repaired by Adenine base at 5′ and then adapters were ligated. Finally, the target cDNA library was amplified by PCR. Validation of the genomic cDNA library was carried out by quantifying and evaluating the quality using Agilent 2100 Bioanalyzer and Qubit respectively. Then the amplified genomic libraries of the fourteen samples were pooled and sequenced with Illumina MiSeq platform using its kit (www.illumina.com).

### Sequence data analysis

The quality of raw sequence reads for each sample was evaluated with fast x-toolkit (hannonlab.cshi.edu) and sequence lead adapters were clipped. Then reads were trimmed at the ends using a dynamic trim program with a quality cut off score limit 0.05 and discarding all read less than 25 bp.

The cleaned reads were assembled following de novo approaches by Trinity assembler using CLC Genomics Workbench 4.5 (CLC Bio, Denmark) with minimum length of 500 bp, mismatch cost two, deletion cost three and insertion cost three. After assembling the reads, sets of contigs were generated and the quality of the assembler was evaluated by assembly statistics value like total trinity transcript, N 50, and average length. Contigs were sorted by size using Solexa QA software and the longest contigs were used for searching similarity from genome sequence database by BLAST N search [30] and the best hit for each query was recorded. After identifying the closest species, the contigs were mapped to reference the viral genome sequence from NCBI database. Comparisons were made using the fragment representing the coat protein gene of the detected viral species.

Coat protein gene sequences of the viruses detected in the present study were deposited in the GenBank with the respective accession numbers (GarV-C, MK336961-64; GarV-D, MK336965-67; GarV-X, MK336968-71; OYDV, MK336972-77; LYSV, MK336978-83).

### Phylogenetic trees

Pairwise molecular diversity for coat protein (CP) nucleotide and amino acid sequences were calculated in MEGA 5.0 [31] using the nucleotide substitution model. Phylogenetic analyses (Maximum Likelihood) of CP nucleotide sequence data of Allexivirus and Potyvirus were conducted using the software package PhyML 3.1 [32]. For phylogenetic analysis, TPM2uf+G and GTR+G models were selected by jmodel test 2.1.5 [33] for the CP nucleotide sequence data of Allexivirus and Potyvirus, respectively. The sequence of PepMV, (Pepino mosaic virus, family Alphaflexiviridae, genus Potexvirus) is used as the outgroup for Allexivirus and RMV, (Ryegrass mosaic virus, family Potyviridae, genus Rymovirus) is used as the outgroup for Potyvirus.

## Results

### Percentage occurrence and multiple infections of garlic viruses

The result revealed that five different viral species were isolated from the diseased samples obtained from the Oromia and Amhara regions of Ethiopia. All the garlic samples were infected by mixtures of different viral species. The viruses that belong to Allexivirus and Potyvirus were found to be the most common viral species in all tested garlic samples. OYDV, LYSV, Garlic virus C (GarV-C), Garlic virus X (GarV-X), and Garlic virus D (GarV-D) were identified at 100, 92, 92, 92, and 78.6 %, respectively. The three improved varieties are infected by all the virus isolates, mixed infection. Sixty-five percent of the samples had multiple infections with OYDV, LYSV, Garlic virus C, Garlic virus D, and Garlic virus X (Table 2).

**Table 2.** List of identified Garlic viruses, species complex and percentage occurrence

| No | Sample id | Types of viruses detected |  |  |  |  |
| --- | --- | --- | --- | --- | --- | --- |
|  |  | OYDV | LYSV | Gar VC | Gar VD | Gar VX |
| 1. | 6 | ✓ | ✓ | ✓ | ✓ | ✓ |
| 2. | 9 | ✓ | x | ✓ | ✓ | ✓ |
| 3. | 10 | ✓ | ✓ | ✓ | ✓ | ✓ |
| 4. | 23 | ✓ | ✓ | ✓ | x | ✓ |
| 5. | 35 | ✓ | ✓ | ✓ | ✓ | ✓ |
| 6. | 55 | ✓ | ✓ | ✓ | ✓ | ✓ |
| 7. | 80 | ✓ | ✓ | ✓ | ✓ | X |
| 8. | 98 | ✓ | ✓ | ✓ | x | ✓ |
| 9. | 102 | ✓ | ✓ | x | x | ✓ |
| 10. | 138 | ✓ | ✓ | ✓ | ✓ | ✓ |
| 11. | 153 | ✓ | ✓ | ✓ | ✓ | ✓ |
| 12. | BN* | ✓ | ✓ | ✓ | ✓ | ✓ |
| 13. | TS* | ✓ | ✓ | ✓ | ✓ | ✓ |
| 14. | V3* | ✓ | ✓ | ✓ | ✓ | ✓ |
| % of occurrence |  | 100 | 92 | 92 | 78 | 92 |

OYDV, Onion yellow dwarf virus; LYSV, Leek yellow stripe virus; Gar VC, *Garlic virus C*; Gar VD, *Garlic virus D*; Gar VX, *Garlic virus X*
\*BN, Bishoftu nech; TS, Tseday; V3, kuriftu

### Allexivirus evolutionary divergence

Allexivirus group namely Garlic virus C (GarV-C) was one of the most prevalent viral species with four characterized isolated strains based on the CP gene. The isolated strains had shown nucleotide sequence identity with the closest type strains in the range of 81.2-91.8 % (Table 3). Isolated strain S24GVC had 91.8% nucleotide sequence identity with the closest type reference strain JX488637, AB010302, and JX488621; similarly, strain S3GVC and S1GVC had nucleotide identity of 91.8% and 91.4% with the type strain AB010302, respectively. Among the isolated strains, strain S9GVC had shown the lowest nucleotide sequence identity of 81.2%, 81.9%, 81.9, and 82.9% with the type strains KF95556, HM777004, KX034776, and JX488640, respectively.

**Table 3.** Pairwise nucleotide and amino acid sequence identity for *Garlic virus C*

| Accession No | % Nucleotide Sequence Identity |  |  |  | % Amino Acid Sequence Identity |  |  |  |
| --- | --- | --- | --- | --- | --- | --- | --- | --- |
|  | S1GVC | S24GVC | S3GVC | S9GVC | S1GVC | S24GVC | S3GVC | S9GVC |
| **JX488621** | 91 | 91.8 | 91.4 | 86 | 95.6 | 94.7 | 96.1 | 92.5 |
| **JX488644** | 90.8 | 91.7 | 91.2 | 86 | 95.6 | 94.7 | 96.1 | 92.5 |
| **JX488637** | 91 | 91.8 | 91.4 | 86 | 95.6 | 94.7 | 96.1 | 92.5 |
| **JQ899448** | 90.3 | 90.4 | 90.7 | 85.5 | 95.6 | 94.7 | 96.1 | 92.5 |
| **AB010302** | 91.4 | 91.8 | 91.8 | 86.3 | 96.1 | 95.2 | 96.5 | 93 |
| **KX034775** | 90.3 | 89.1 | 90.7 | 83.9 | 96.1 | 95.6 | 96.5 | 93 |
| **HM777004** | 89.7 | 86.9 | 90.1 | 81.8 | 94.7 | 94.3 | 95.2 | 91.7 |
| **KF955566** | 90 | 87.4 | 90.4 | 81.2 | 94.7 | 94.3 | 95.2 | 91.7 |
| **JX488640** | 90.5 | 89 | 91.2 | 82.9 | 93.9 | 93 | 94.3 | 90.4 |
| **KX034776** | 89.4 | 88.3 | 90.1 | 81.9 | 94.7 | 95.2 | 95.2 | 91.2 |

The amino acid sequence data of isolated strains under GarV-C species revealed a 90.4 - 95.6% sequence identity as compared with the closest type reference strains (Table 3). Strain S3GVC had 96.5% and 96.1% amino acid sequence similarity with the type reference strain AB010302 and KX034775; and JX488621, JX488644, JX488637, and JQ899448, respectively. While strain S24GVC and S1GVC had a 95.6% amino acid sequence identity with JX488621, JX488644, JX488637, and KX034775, respectively. Among the isolated strains, strain S9GVC had shown the lowest amino acid sequence identity of 90.4% with the type reference strain JX488640.

Allexivirus strain identified under Garlic virus D (GarV-D) revealed 94.2 - 98.3% nucleotide sequence identity with the closest type strains (Table 4). Isolated strain S22GVD had shown 98.3% and 98.2%nucleotide sequence identity with that of the type reference strains KX889779 and KX889779; and KX889778, respectively. While strain S3GVD had shown 97.9% and 97.5% nucleotide sequence identity with the type reference strains AB010303 and KF555653%; and KF446188%, respectively. Relatively the lowest nucleotide divergence in this group was obtained by strain S22GVD with 94.2% and 94.6% nucleotide sequence identity with the type reference strain KF446207 and AB010303, respectively. The percentage CP amino acid sequence identity of isolated strains in Garlic virus D was ranged from 99.6 - 95.2% (Table 4). The highest amino acid sequence identity of 99.6% was obtained by strain S23GVD as compared with the type reference strains KX889778, KX889780, KF446194, and KX889779. While strains S3GVD and S22GVD had shown 97.4 - 98.2% and 95.2 - 96.9% amino acid sequence identity with the closest type reference strain.

**Table 4.**
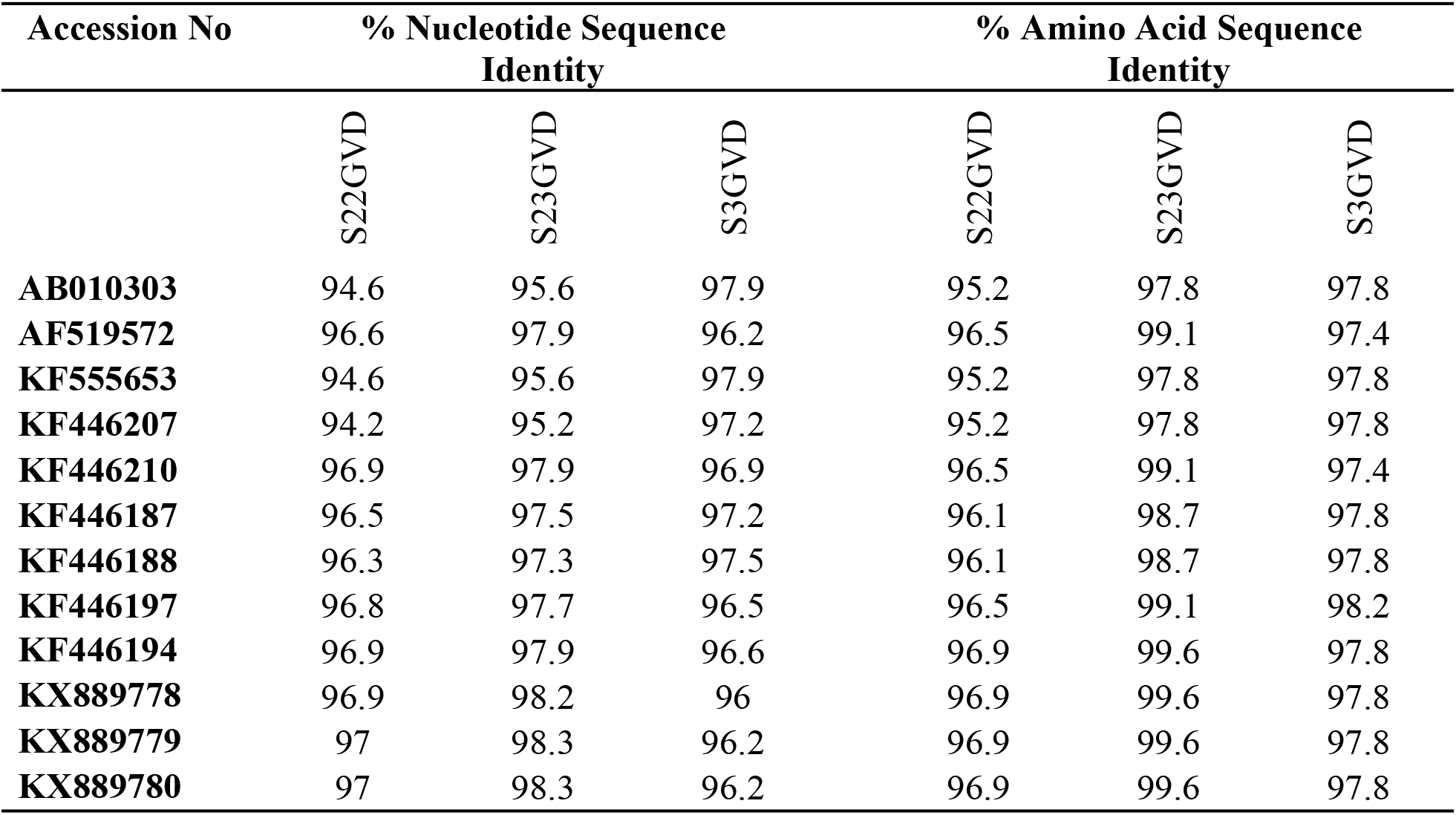
Pairwise nucleotide and amino acid sequence identity for *Garlic virus D*

The percentage nucleotide identity values for the isolated Allexivirus CP gene of Garlic virus X (GarV-X) ranged from 92.8 - 98.4% (Table 5). The isolated strain S12GVX and S9GVX had 98.4% coat protein (CP) nucleotide sequence identity with the closest type strain JQ807994 and KF530328, followed by strain S7GVX which had a sequence identity of 97.9% with the type strain JQ807994. While strain S24GVX had 96.3% nucleotide sequence identity with the type strains JQ807994, KF471313, and KF530328 (Table 5). On the other hand, the amino acid sequence identity of the isolated Allexivirus strains in the GarV-X were ranged from 100 - 96.9%. Strain S12GVX, S9GVX, and S7GVX had 100% amino acid sequence similarity with the closest type reference strain KF471313 and KF530328, while the isolated strain S24GVX had 98.2% amino acid sequence similarity with the above type reference strains (Table 5).

**Table 5.** Pairwise nucleotide and amino acid sequence identity for *Garlic virus X*

| Accession No | % Nucleotide Sequence Identity |  |  |  | % Amino Acid Sequence Identity |  |  |  |
| --- | --- | --- | --- | --- | --- | --- | --- | --- |
|  | S12GVX | S24GVX | S9GVX | S7GVX | S12GVX | S24GVX | S9GVX | S7GVX |
| **GQ475426** | 94.8 | 93.2 | 94.8 | 94.5 | 98.2 | 97.4 | 98.2 | 98.2 |
| **JQ807994** | 98.4 | 96.3 | 98.4 | 97.9 | 99.1 | 97.4 | 99.1 | 99.1 |
| **KF471313** | 98.3 | 96.3 | 98.3 | 97.7 | 100 | 98.2 | 100 | 100 |
| **KF530328** | 98.3 | 96.3 | 98.3 | 97.7 | 100 | 98.2 | 100 | 100 |
| **LN875276** | 94.8 | 92.8 | 94.8 | 94.2 | 99.1 | 97.4 | 99.1 | 99.1 |
| **LN875277** | 94.9 | 92.9 | 94.9 | 94.4 | 99.1 | 97.4 | 99.1 | 99.1 |
| **HQ873847** | 94.6 | 93.1 | 94.6 | 94.4 | 98.2 | 97.4 | 98.2 | 98.2 |
| **JX429969** | 95.5 | 94.1 | 95.5 | 95.2 | 98.7 | 96.9 | 98.7 | 98.7 |

### Potyvirus evolutionary divergence

The percentage nucleotide identity values for the isolated Potyvirus CP gene of Onion yellow dwarf virus (OYDV) ranged from 83.3 - 97.6% (Table 6). The largest CP nucleotide identity was observed among a pairwise comparison of the isolated strain S23OYDV versus with that of the type reference strain GQ475394 (97.6%), GQ475393 (97.4%), GQ475396 and GQ475397 (96.4%), and GQ475405 (96.2%) followed by 92.3% CP nucleotide identity among isolated strain S21OYDV versus AJ292231 and AJ510223. For strain S22OYDV, S12OYDV, S1OYDV, and S20OYDV the closest relative reference strains for CP nucleotide identity were observed with AJ292231 and AJ510223 (88.3%); AJ409311 (88.3%); KF862684 (87.4%); AJ409311 (87.6%), respectively. Relatively the lowest pairwise nucleotide identity of 83.3% was observed by strain S20OYDV as compared with the reference strains GQ475393, GQ475398, and GQ475397.

**Table 6.** Pairwise nucleotide and amino acid sequence identity for *Onion yellow dwarf virus* (OYDV)

| Accession No | % Nucleotide Sequence Identity |  |  |  |  |  | % Amino Acid Sequence Identity |  |  |  |  |  |
| --- | --- | --- | --- | --- | --- | --- | --- | --- | --- | --- | --- | --- |
|  | S1OYDV | S12OYDV | S20OYDV | S23OYDV | S21OYDV | S22OYDV | S1OYDV | S12OYDV | S20OYDV | S23OYDV | S21OYDV | S22OYDV |
| **AJ409311** | 87.4 | 88.3 | 87.6 | 84.9 | 85.6 | 85.7 | 93.6 | 92.8 | 92.8 | 88.6 | 90.3 | 89.8 |
| **KF862684** | 87.7 | 86.5 | 87.2 | 85.3 | 84.8 | 86.1 | 93.6 | 92.4 | 93.2 | 90.3 | 89.8 | 90.3 |
| **GQ475358** | 87 | 87.6 | 86.8 | 84.6 | 85.7 | 84.8 | 91.9 | 91.1 | 91.9 | 88.1 | 89 | 88.1 |
| **KF862685** | 86.9 | 88 | 87.4 | 84.8 | 85.7 | 85 | 92.4 | 92.4 | 93.2 | 88.6 | 89.8 | 89.4 |
| **GQ475360** | 86.9 | 87.6 | 87.2 | 84.5 | 84.9 | 85 | 92.8 | 91.9 | 92.8 | 88.1 | 89.4 | 89 |
| **GQ475359** | 86.5 | 87 | 86.5 | 84.2 | 84.9 | 84.5 | 91.9 | 91.1 | 91.9 | 87.7 | 89 | 88.1 |
| **AJ510223** | 84.1 | 86.1 | 84.9 | 88.5 | 92.3 | 88.3 | 88.6 | 91.1 | 90.7 | 95.3 | 94.5 | 92.4 |
| **AJ292231** | 84.1 | 86.1 | 84.9 | 88.5 | 92.3 | 88.3 | 88.6 | 91.1 | 90.7 | 95.3 | 94.5 | 92.4 |
| **AJ409309** | 84.1 | 83.7 | 83.4 | 87.6 | 88.7 | 86.5 | 88.1 | 89.8 | 89.4 | 92.8 | 92.8 | 91.1 |
| **AJ307033** | 85 | 85.3 | 83.8 | 88.8 | 89.9 | 87 | 88.6 | 90.3 | 89.8 | 92.8 | 93.6 | 91.1 |
| **KF632714** | 85.3 | 85.7 | 85.7 | 86.6 | 89.5 | 86.8 | 91.5 | 91.1 | 91.1 | 93.2 | 91.5 | 91.9 |
| **GQ475394** | 85.2 | 83.8 | 83.4 | 97.6 | 84.6 | 84.1 | 89 | 89 | 88.6 | 97.9 | 89 | 88.6 |
| **GQ475393** | 85 | 83.7 | 83.3 | 97.4 | 84.5 | 83.9 | 88.6 | 88.6 | 88.1 | 97.5 | 88.6 | 88.1 |
| **GQ475398** | 85.8 | 84.1 | 83.3 | 96.4 | 84.5 | 84.5 | 89.4 | 89.4 | 89 | 97.5 | 89.4 | 89 |
| **GQ475405** | 85.7 | 84.2 | 83.4 | 96.2 | 84.3 | 84.3 | 89.4 | 89.4 | 89 | 97.5 | 89.4 | 89 |
| **GQ475397** | 85.8 | 84.1 | 83.3 | 96.4 | 84.5 | 84.5 | 89.4 | 89.4 | 89 | 97.5 | 89.4 | 89 |

The result of CP amino acid identity across all the isolated strains of garlic OYDV versus with that of the type of reference strains was ranged from 88.1 - 97.9% (Table 6). Isolated strain S23OYDV had the maximum CP amino acid identity of 97.9%, 97.5%, and 95.3% with that of the type reference strain GQ475394; GQ475393, GQ475398, GQ475405 and GQ475397; and AJ510223 and AJ292231, respectively. The other closest relative type reference strains AJ292231 and AJ510223 had CP amino acid identity of 94.5% with the isolated strain S21GOYDV, while strains S1OYDV, S20OYDV, S12OYDV, and S22OYDV had shown CP amino acid sequence identity with AJ409311(93.6%); KF862684 and KF862685 (93.2%); AJ409311 (92.8%); and AJ510223 and AJ292231 (92.4%), respectively. Relatively the lowest pairwise amino acid sequence identity of 88.1% was observed by strain S1OYDV, S20OYDV, S23OYDV, and S22OYDV as compared with the reference strain AJ409309; GQ475393; GQ475360; GQ475359 and GQ475393, respectively (Table 6).

Pairwise comparison of isolated garlic Leek yellow stripe virus (LYSV) showed nucleotide and amino acid CP sequence identity with that of the type reference strain in a range of 81.8 - 96.1% and 81.4 - 96.2%, respectively (Table 7). The comparison showed that isolated strain S14GVY had 96.1% and 96.2% CP nucleotide and amino acid identity with that of the type reference strain AJ307057 and AJ292225, respectively. While strain S15GVY had 90.3% CP nucleotide and 95.3% CP amino acid sequence identity with the above type reference strain. The other strain including S1GVY, S20GVY, S23GVY, and S4GVY had shown CP nucleotide identity 81.8 - 91.1% with that of the closest type reference strain. Among the isolated strains, strain S4GVY and S1GVY showed the lowest nucleotide sequence identity of 81.8% and 82.1% with the reference strains AB194641 and DQ296002, respectively. Among the isolated strains, strain S1GVY showed the lowest amino acid sequence identity of 81.8% with the reference strains AB194642, DQ296002, AB194641, and DQ402056 (Table 7).

**Table 7.** Pairwise nucleotide and amino acid sequence identity for *Leek yellow stripe virus* (LYSV)

| Accession No | % Nucleotide Sequence Identity |  |  |  |  |  | % Amino Acid Sequence Identity |  |  |  |  |  |
| --- | --- | --- | --- | --- | --- | --- | --- | --- | --- | --- | --- | --- |
|  | S15LYSV | S1LYSV | S14LYSV | S20LYSV | S23LYSV | S4LYSV | S15LYSV | S1LYSV | S14LYSV | S20LYSV | S23LYSV | S4LYSV |
| AJ307057 | 90.3 | 85.8 | 96.1 | 88.9 | 87.9 | 88.8 | 95.3 | 87.7 | 96.2 | 92.8 | 89.8 | 91.1 |
| AJ292225 | 90.3 | 85.8 | 96.1 | 88.9 | 87.9 | 88.8 | 95.3 | 87.7 | 96.2 | 92.8 | 89.8 | 91.1 |
| AB194635 | 88.3 | 85.3 | 91.2 | 86.6 | 85.4 | 85.6 | 94.9 | 88.6 | 94.1 | 91.9 | 90.7 | 91.1 |
| AB194634 | 88.1 | 85.2 | 90.8 | 86.2 | 85.3 | 85.2 | 94.9 | 88.6 | 94.1 | 91.9 | 90.7 | 91.1 |
| AJ409305 | 87.0 | 83.9 | 89.5 | 84.6 | 84.8 | 85.2 | 92.8 | 86.4 | 92.8 | 89.8 | 89.0 | 89.8 |
| DQ299382 | 86.9 | 83.7 | 86.9 | 84.2 | 86.8 | 84.8 | 91.5 | 86.4 | 91.5 | 89.0 | 88.6 | 89.0 |
| KF597283 | 86.9 | 83.7 | 86.9 | 84.2 | 86.8 | 84.8 | 91.9 | 86.9 | 91.9 | 89.4 | 89.0 | 89.4 |
| JN127339 | 86.5 | 83.7 | 87.4 | 84.6 | 85.2 | 84.5 | 93.6 | 87.7 | 93.2 | 91.5 | 89.8 | 90.3 |
| AF538950 | 85.3 | 82.2 | 85.4 | 83.3 | 85.7 | 82.9 | 91.5 | 85.6 | 90.7 | 89.0 | 89.4 | 88.1 |
| AF314146 | 85.6 | 83.4 | 88.7 | 84.9 | 85.4 | 84.5 | 91.9 | 86.4 | 91.1 | 89.4 | 88.6 | 88.6 |
| AB005611 | 85.8 | 82.6 | 86.0 | 83.3 | 85.3 | 82.7 | 88.6 | 82.6 | 87.3 | 86.4 | 86.9 | 84.3 |
| AB194642 | 85.8 | 82.6 | 85.8 | 83.1 | 85.3 | 82.7 | 87.7 | 81.8 | 86.4 | 85.6 | 86.0 | 83.5 |
| DQ296002 | 85.6 | 82.1 | 85.3 | 82.6 | 85.2 | 81.9 | 87.7 | 81.8 | 86.4 | 85.6 | 86.0 | 83.5 |
| AB194638 | 84.8 | 82.2 | 85.6 | 83.3 | 84.1 | 83.3 | 90.7 | 86.0 | 89.0 | 88.1 | 89.4 | 88.1 |
| AB194641 | 85.4 | 82.2 | 84.9 | 82.5 | 84.6 | 81.8 | 87.3 | 81.4 | 86.0 | 85.2 | 85.6 | 83.1 |
| DQ402056 | 85.8 | 82.6 | 85.8 | 83.1 | 85.3 | 82.5 | 87.3 | 81.4 | 86.0 | 85.2 | 85.6 | 83.1 |
| AJ409308 | 85.3 | 82.6 | 88.4 | 84.5 | 84.9 | 83.0 | 89.8 | 83.9 | 89.0 | 88.1 | 87.7 | 86.0 |

### Phylogenetic tree for Allexivirus and Potyvirus

Phylogenetic analysis of CP nucleotide sequences confirmed the taxonomic placement of three Allexivirus isolates identified on garlic from Ethiopia as members of GarV-C, GarV-D and GarV-X. Ethiopian Isolates of GarV-C namely S1GVC, S3GVC, and S24GVC were clustered with isolates of Brazil, Australia, Japan, Poland, and Argentina (Fig 1), while S9GVC of Ethiopian isolate was the most divergent among the group. The other isolates identified as members of GarV-D such as S3GVD, S22GVD, and S23GVD were clustered and closely related to isolates from Argentina, Japan, Poland, and Korea (Fig 1). Isolates identified in the members of GarV-X including S7GVX, S9GVX, S12GVX, and S24GVX were clustered with isolates from Australia, Brazil, Hungary, Spain, Italy, and China (Fig 1).

**Fig 1.**
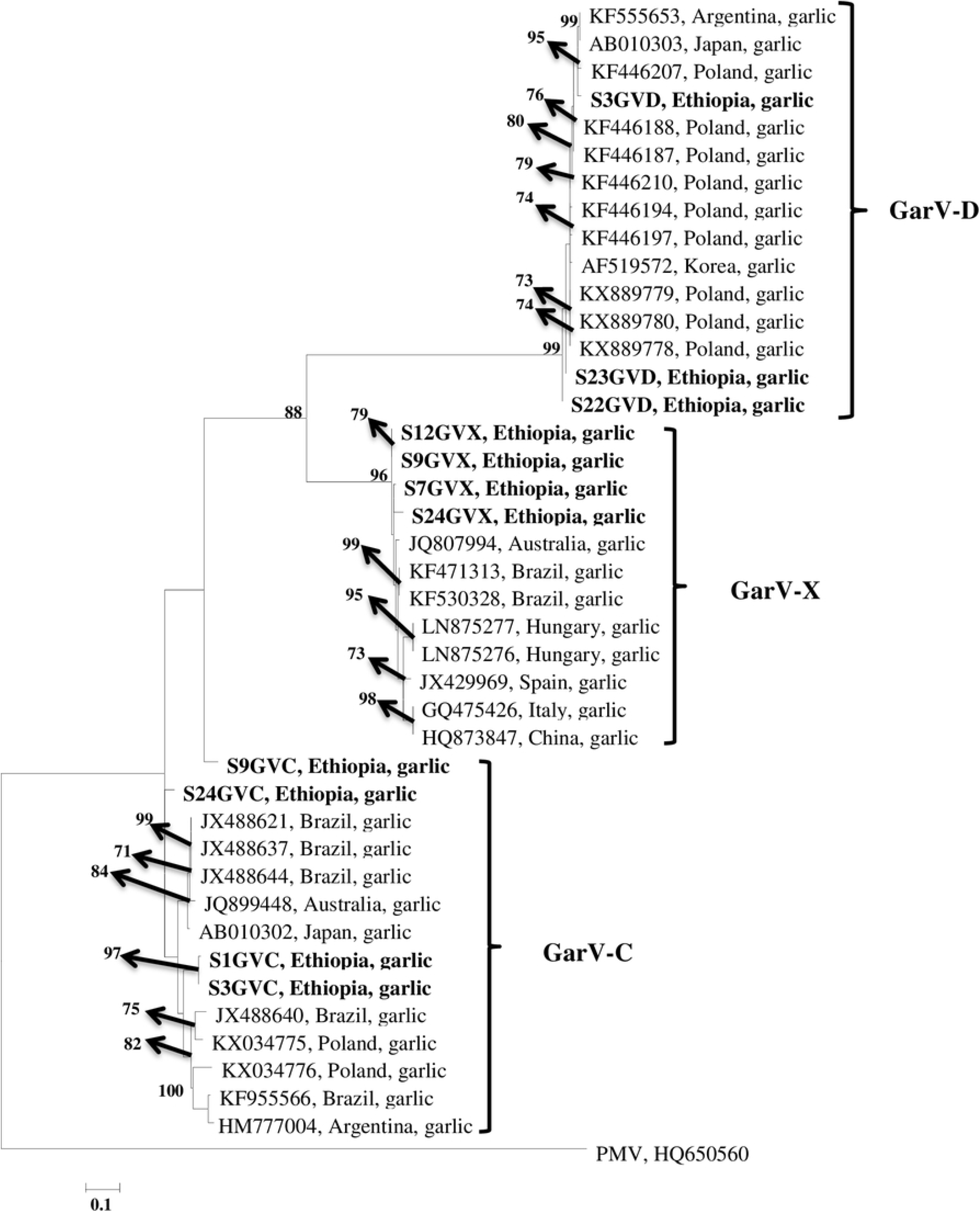
Allexivirus isolates identified in the study. Phylogenetic analysis of Coat protein (CP) nucleotide sequences of Allexivirus inferred by Maximum likelihood using the program PhyML. Branches were supported by bootstrap values greater than 70% (based on 100 replicates). Pepino mosaic virus (PMV) is provided as the outgroup.

Phylogenetic analysis of CP nucleotide sequences of 42 OYDV isolates containing 6 Ethiopian and 36 other isolates from different parts of the world separated those isolates from Ethiopia into two groups. The only isolate, S23OYDV, from Ethiopian samples in this study, separated from the rest and was clustered and closely related to OYDV isolates sourced from garlic representing Italy (GQ475393, GQ475394, GQ475397, GQ475398, and GQ475405) (Fig 2). The other isolates S21OYDV and S21OYDV were clustered with isolates sourced from garlic from China (AJ292231, AJ307033, and AJ510223) and Argentina (KF632714), while the rest three isolates S1OYDV, S12OYDV, and S20OYDV were clustered with isolates all from garlic from Poland (KF862685), USA (GQ475358 and GQ475360) and China (AJ409311). These isolates were very closely related to OYDV representing Poland, the USA, and China (Fig 2). While Phylogenetic analysis of CP nucleotide sequences of 44 LYSV isolates containing 6 Ethiopian and 38 other isolates from around the world separated those isolates from Ethiopia into one group. Ethiopian isolates were clustered with isolates of China (AJ292225, AJ307057, and AJ409305) and Korea (AB194634 and AB194635). The result revealed that Ethiopian isolates in this study are very closely related to LYSV isolates from garlic representing China and Korea (Fig 3).

**Fig 2.**
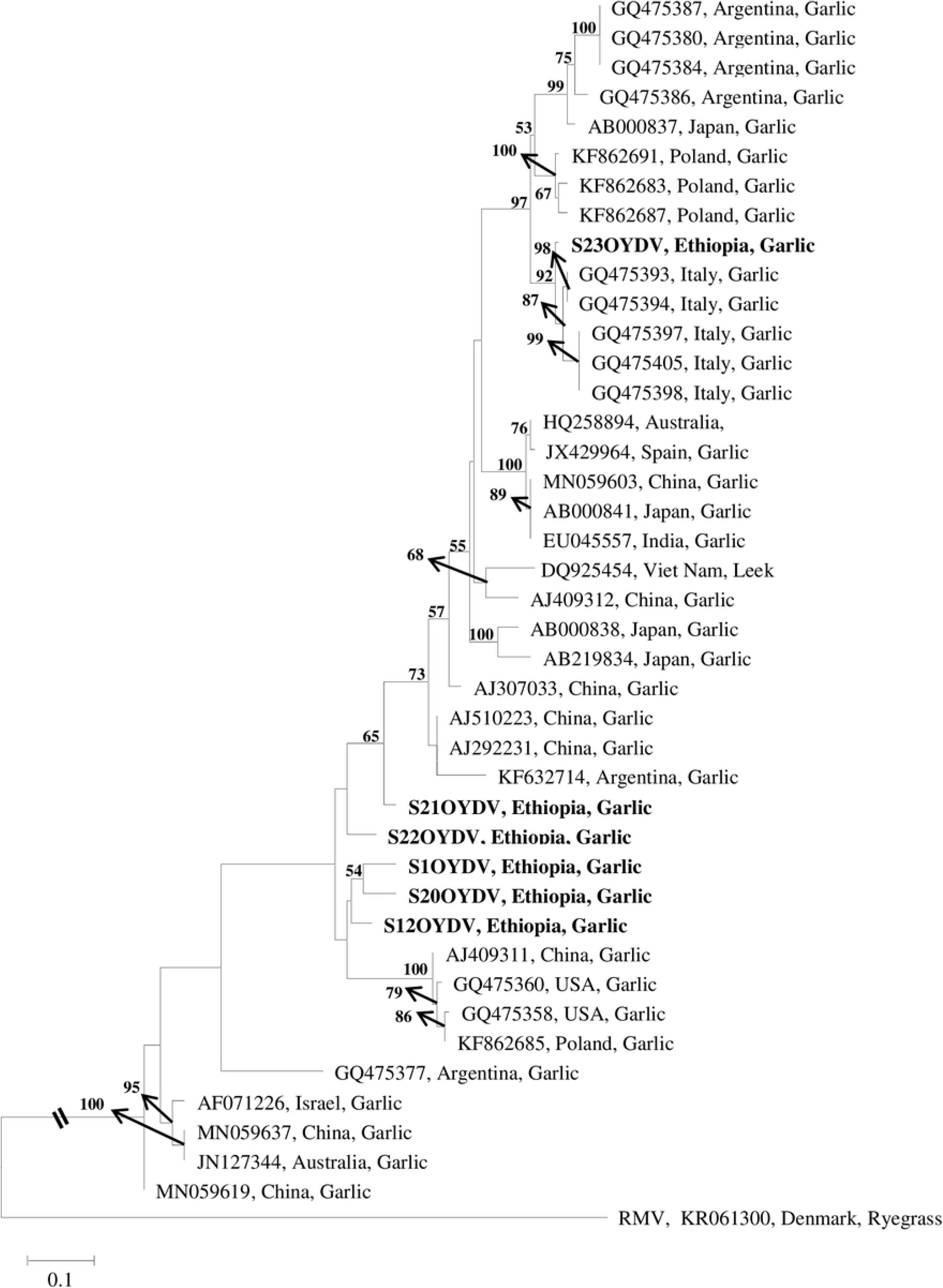
Onion yellow dwarf virus isolates. Phylogenetic analysis of Coat protein (CP) nucleotide sequences of Onion Yellow Dwarf Virus (OYDV) inferred by Maximum likelihood using the program PhyML. Branches were supported by bootstrap values greater than 50% (based on 100 replicates). Ryegrass mosaic virus (RMV) is provided as the outgroup.

**Fig 3.**
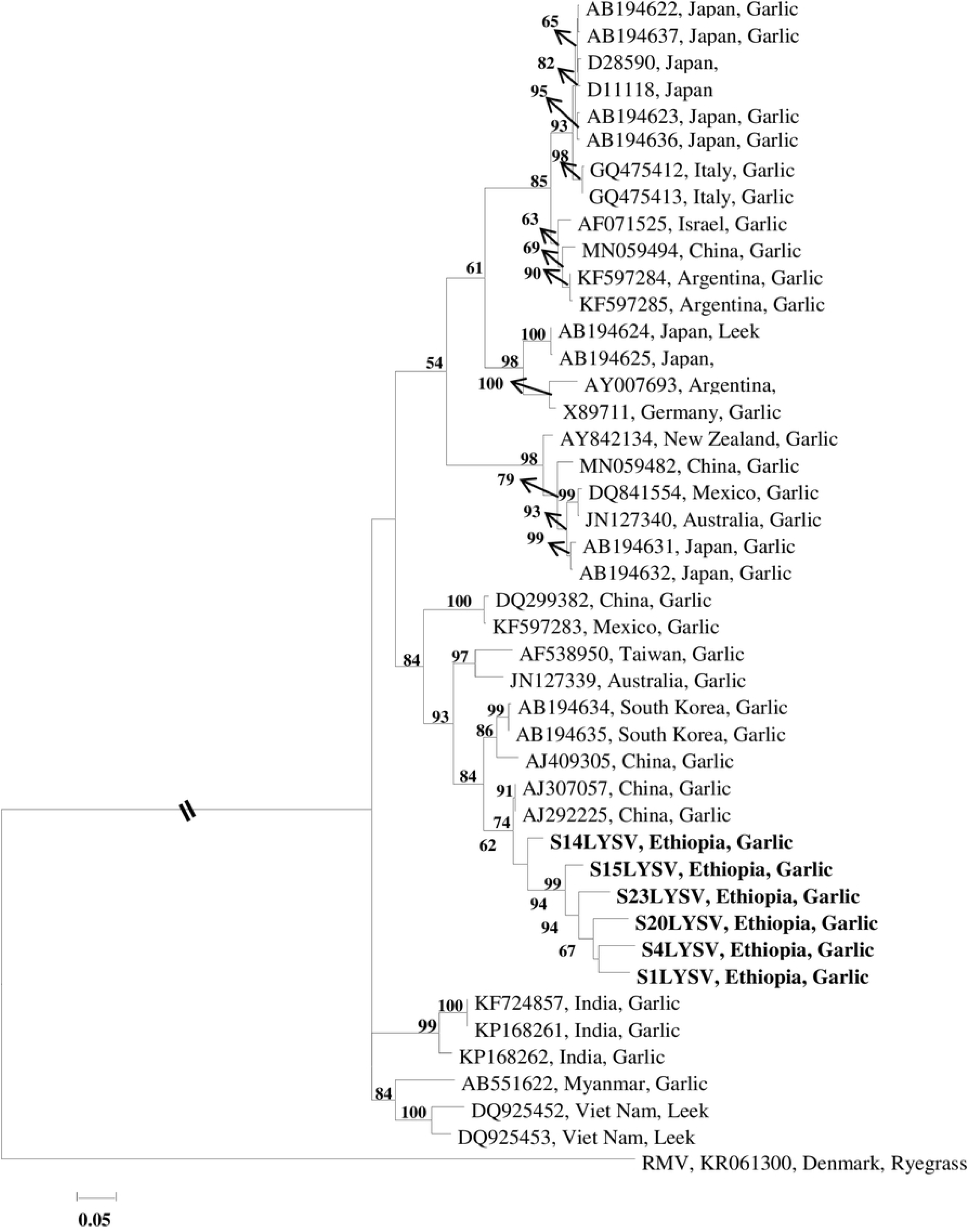
Leek yellow stripe virus isolates. Phylogenetic analysis of Coat protein (CP) nucleotide sequences of Leek Yellow Stripe Virus (LYSV) inferred by Maximum likelihood using the program PhyML. Branches were supported by bootstrap values greater than 50% (based on 100 replicates). Ryegrass mosaic virus (RMV) is provided as the outgroup.

## Discussion

Ethiopia is one of the ten garlic producer countries where garlic production has been seriously challenged by viral diseases. This study is therefore aimed to identify garlic infecting viruses and its complex occurrence by targeting coat protein gene sequences. The results from the study showed that five viruses identified viz. Onion yellow dwarf virus and Leek yellow stripe virus from Potyvirus and Garlic virus C, Garlic virus D, and Garlic virus X from Allexivirus. In all tested samples co-infection of Potyvirus and Allexivirus were identified. This multiple distribution and co-occurrence of the different garlic viral species complex of this study suggested that the viruses have mostly been distributed through infected bulbs. In Ethiopia, the distribution of virus-free garlic planting materials among farmers is truly uncommon. Market places are the most common sources of planting materials that contribute to the occurrence of multiple infections of the virus across the country since the bulbs are getting into the market from different parts of the country. In most cases, the leftover bulbs from consumption were also used as sources of planting materials. Such cases of multiple occurrences of garlic viral complex in a production field were strongly suggested by several other authors around the globe (Bereda et al. 2017; Stephen et al. 2012; Sward and Brennan 1994).

OYDV was the most prevalent followed by LYSV, Garlic virus X and Garlic virus C, and Garlic virus D in respective order. The study revealed the presence of complex mixtures of viruses from the two genera. Similar results had been reported by several authors (Fayad-Andre et al. 2011; Shahraeen et al. 2008; Shiboleth et al. 2001; Takaichi et al. 1998). In particular, those most prevalent viruses namely OYDV and LYSV which incurred the greatest economic losses were detected in all examined samples in this study, took its priority supported by much research (Bagi et al. 2012; Lunello et al. 2007; Lot et al. 1998). The frequent infection of garlic with Potyviruses associated with the effective transmission capacity of its vector, aphid (Melo et al. 2004), suggested that the on-time disease management scheme needs to be implemented to minimize the devastating synergetic effect of Potyviruses. In support of this study, mosaic symptoms which are the most common phenomena in this study had been identified as the major causes for yield loss and quality (Conci et al. 2010; Cafrune et al. 2006; Conci et al. 2003; Chen et al. 2001).

The coat protein sequence data of the garlic virus generated in this study has proved useful in identifying garlic Allexiviruses and Potyviruses. The pairwise comparison of genetic relatedness for the detected isolates confirmed the presence of complex viral mixture as reported elsewhere (Fayad-Andre et al. 2011; Shahraeen et al. 2008; Takaichi et al. 1998). The study revealed a range of variation in the CP nucleotide sequence data of Allexivirus. Pairwise comparisons of nucleotide sequence data of Allexivirus CP genes gave reliable identification of the isolates. The minimum identity of the CP nucleotide sequences of the detected GarV-C, GarV-D, and GarV-X isolates of this study in comparison with the known members of the species deposited in the GenBank was 81.2%, 94.2%, and 92.8%, respectively. Similarly, the minimum identity of the CP amino acid sequences of the above detected Allexivirus species was 91.2%, 95.2% and 96.9%, respectively. These values revealed that the detected isolates of Allexivirus had high genetic relatedness with the previously identified species that all falling above the species demarcation values of 72% for nucleotide and 80% for the amino acid sequence (Adams et al. 2004). However, regarding the species GarV-C, the identity of the CP nucleotide sequence data of isolate S9GVC with the closest type strains in the Gene Bank was below the minimum value for within species identity (Adams et al. 2004).Whereas, the corresponding pairwise comparisons of amino acid sequence data analyzed within species were relatively in agreement with the within species threshold (Adams et al. 2004). Overall isolates detected in GarV-C species showed divergence with each other and the type reference strains of HM777004, KF955566, JX488640, and KX034776. This is an exception in this study that needs reconsidering the GarV-C complex according to the present within species threshold in particular but not exclusively for the CP nucleotide sequence data.

Pairwise comparisons of the isolated Potyviruses to examine their genetic relatedness with that of the known members of the species indicated the very high identity of 83.3 - 97.6% for OYDV and 81.8 - 96.1% for LYSV. This figure fully supports the high identity of the detected isolates with the closest relatives since the figures are above the species demarcation value of 78 % nucleotide identity [39]. The coat protein amino acid sequence data of Potyviruses were compared with the known members of the species. This comparison revealed the high identity of isolates of 88.1 - 97.9% for OYDV and 81.4 - 96.2% for LYSV. This high value represents a comparison between different isolates of OYDV and LYSV of the same species due to the value above species discrimination of 82 % [39] amino acid sequence data of CP. One exceptional strain is S1LYSV that showed less amino acid identity of 81.4% with reference strains of AB194642, DQ296002, AB194641, and DQ402056.

The lower sequence divergence in the viral CP of LYSV isolates of Ethiopian samples suggested that the virus constitutes a new cluster that probably developed more recently. Similar results were obtained on OYDV isolates with the exception that one isolate S23OYDV was clustered separately as compared with the others. This implies that the clustering of Ethiopian OYDV isolates into two distant clades indicate that the virus has been relatively well established in Ethiopia as compared with the LYSV isolates. Hence, as suggested by García-Arenal et al. (2001) the relatively higher values of the genetic diversity among sequences of the two clusters on OYDV isolates can indicate older populations of OYDV as compared with LYSV isolates of this study.

Phylogenetic analysis of Allexiviruses revealed isolates in this genus showed strong bootstrap value support for each species in the clad. In particular, isolate S9GVC in the GarV-C clade showed a high divergence from isolates of the same species in the phylogenetic tree where it belongs. The isolate S9GVC could not grouped in any clade, showing the presence of polytomy and indicating a lack of data that could point to the origin of these sequences as suggested by (Vučurović et al. 2017). Hence, the existing variation pattern complexity in this isolate suggested the isolate may have had different evolutionary processes.

The phylogenetic analysis of Potyvirus also revealed no correlation between geographic localization and CP sequence variation in LYSV and OYDV of this study and other CP sequences obtained from the rest of the world. The study suggested that the frequently mixed infection observed in this study between different genotypes of the same species in a single plant and the vegetative propagation nature of garlic might be the cause of the high variability of the garlic viruses. Furthermore, the result represents the presence of independent influences of different evolutionary forces of geographically distant virus isolates and/or selection pressure for the adaptation of different geographical localization (Koo et al. 2002; Fajardo et al. 2001).

## Conclusion

The study presents and concludes that garlic viral species in the genus Potyvirus and Allexivirus are identified. The study also indicated the presence of garlic viral species complex in the production fields of Ethiopia. The analysis of nucleotide and amino acid sequences of coat protein genes revealed three species from Allexivirus and two from Potyvirus in complex mixtures. However, there might have been the possibility of garlic plant infection by more other viruses which is not yet detected in this particular study since there is an uncontrolled exchange of infected planting material among growers in the country. Furthermore, the result explained that the high prevalence and detection of OYDV and LYSV from all garlic experimental samples suggested the association of reduction of garlic yield with these viruses in synergy with the other Allexiviruses. The other finding in this study was the high genetic divergence observed among isolates in the GarV-C species that suggests reconsidering the within species threshold in this group. Finally, our results highlight to set up a working framework to establish virus-free garlic planting material exchange in the country which could result in the reduction of viral gene flow across the country.

## Acknowledgments

The author wishes to thank Debre-Zeit Agricultural Research Center for providing the experimental materials and Bioscience Eastern and Central Africa-International Livestock Research Institute (BecA-ILRI) for the placement and African Bioscience Challenge Fund (ABCF) for the fellowship.

